# Classifying cancer genome aberrations by their mutually exclusive effects on transcription

**DOI:** 10.1101/122549

**Authors:** Jonathan B. Dayton, Stephen R. Piccolo

**Affiliations:** Department of Biology, Brigham Young University, Provo, UT 84602, United States of America; Department of Biomedical Informatics, University of Utah, Salt Lake City, UT 84108, United States of America

**Keywords:** Cancer genomics, integrative omics, machine learning, drug repurposing, pan cancer

## Abstract

**Background:** Malignant tumors are typically caused by a conglomeration of genomic aberrations—including point mutations, small insertions, small deletions, and large copy-number variations. In some cases, specific chemotherapies and targeted drug treatments are effective against tumors that harbor certain genomic aberrations. However, predictive aberrations (biomarkers) have not been identified for many tumor types and treatments. One way to address this problem is to examine the downstream, transcriptional effects of genomic aberrations and to identify characteristic patterns. Even though two tumors harbor different genomic aberrations, the transcriptional effects of those aberrations may be similar. These patterns could be used to inform treatment choices.

**Methods:** We used data from 9300 tumors across 25 cancer types from The Cancer Genome Atlas. We used supervised machine learning to evaluate our ability to distinguish between tumors that had mutually exclusive genomic aberrations in specific genes. An ability to accurately distinguish between tumors with aberrations in these genes suggested that the genes have a relatively different downstream effect on transcription, and vice versa. We compared these findings against prior knowledge about signaling networks and drug responses.

**Results:** Our analysis recapitulates known relationships in cancer pathways and identifies gene pairs known to predict responses to the same treatments. For example, in lung adenocarcinomas, gene-expression profiles from tumors with somatic aberrations in EGFR or MET were negatively correlated with each other, in line with prior knowledge that MET amplification causes resistance to EGFR inhibition. In breast carcinomas, we observed high similarity between PTEN and PIK3CA, which play complementary roles in regulating cellular proliferation. In a pan-cancer analysis, we found that genomic aberrations in BRAF and VHL exhibit downstream effects that are clearly distinct from other genes.

**Conclusion:** We show that transcriptional data offer promise as a way to group genomic aberrations according to their downstream effects, and these groupings recapitulate known relationships. Our approach shows potential to help pharmacologists and clinical trialists narrow the search space for candidate gene/drug associations, including for rare mutations, and for identifying potential drug-repurposing opportunities.

## Background

Typically, a single tumor contains anywhere from tens to millions of genomic aberrations—including point mutations, small insertions, small deletions, and large copy number variations—that differ from the patients normal cells[1–4]. Knowledge of these aberrations may be useful in guiding therapeutic decisions. In some cases, a genomic aberration is the target of an existing therapy and thus may indicate that the therapy is a good match for that patient. For example, *Trastuzumab* is a targeted therapy for HER2-amplified breast cancers[5]. In other cases, a genomic aberration may be a biomarker for an existing therapy, even though the therapy was not explicitly designed to target that aberration[6]. Many such relationships have been identified for combinations of genomic aberration and therapy[7]. However, in many cases, tumors contain no therapeutic biomarker. Furthermore, few gene/drug associations have been made for the “long tail” of genomic aberrations that occur infrequently at the population level[8]. Although it may be economically infeasible to develop targeted therapies for every rare mutation, we may be able to repurpose existing cancer treatments by identifying similarities in tumor biology between tumors that harbor rare and common aberrations.

By disrupting signaling cascades—or pathways—within tumor cells, genomic aberrations can cause the tumor to grow, divide, or dedifferentiate in an uncontrolled manner[9]. Genomic aberrations within tumors are highly variable across cancer patients—each tumor carries a unique panoply of genomic aberrations. However, a much smaller number of signaling cascades is affected. Even though different genes are mutated in two different tumors, these mutations may affect common signaling cascades (e.g., Ras → Raf → MEK → ERK)[10]. We may be able to better understand the effects of genomic aberrations by considering such downstream effects.

Although it is possible to place genomic aberrations in the context of biological pathways, it may be difficult to decipher whether two aberrations have a similar effect on tumor biology, even though they occur within the same signaling cascade. This observation may be especially true for rare mutations, because little is known about the roles they play in tumorigenesis or therapeutic responses, and samples sizes are small. An alternative approach for understanding the effects of these mutations and their potential as biomarkers is to evaluate the transcriptional effects of the mutations. Using existing, high-throughput technologies (e.g., microarrays and RNA-Sequencing), it is possible to quantify gene-expression levels across the entire transcriptome for a modest cost. Two tumors may have similar gene-expression profiles, even though they have no genomic aberrations in common, ostensibly because the aberrations in either tumor have led to similar downstream effects. Therefore, the tumors may respond similarly to drug treatments. When this approach is applied to many tumors, it may be possible to identify transcriptional patterns that can be used as biomarkers of treatment response, independent of the genomic aberrations that occur within these tumors.

We evaluated this idea using publicly available data from The Cancer Genome Atlas (TCGA). We acquired data representing mutations (SNVs, insertions, or deletions), copy-number variations (large amplifications or deletions), and transcription for 9300 tumors across 25 cancer types available in TCGA. We identified tumors that carried mutations in frequently mutated genes (e.g., KRAS, EGFR, and ERBB2) and in genes that are mutated relatively rarely. We made the simplifying assumption that mutations at different genomic loci within a given gene exert a similar effect on tumor biology. We also assumed that mutations in individual genes have a characteristic effect on tumor transcription, despite the presence of additional mutations within each of these tumors.

Initially focusing on lung adenocarcinomas and then extending our analysis to other tumor types, we identified relatively common mutations and filtered the data to include only tumors where these mutations occurred in a mutually exclusive manner—harboring only one of the mutations of interest. Having classified the tumors by mutation status, we used a supervised, machine-learning algorithm to predict mutation status based on transcriptional patterns observed in the tumors. In many cases, the transcriptional patterns were highly predictive of mutation status, thus indicating that individual genomic aberrations influence transcription in distinct ways. Finally, using lower-frequency genes that had been excluded from the initial analysis, we identified genes that, when mutated, resulted in transcriptional patterns that were similar to some of the genes from our initial set. In several cases, these similarities coincided with prior knowledge about cancer pathways as well as with known therapeutic biomarkers.

Our findings show promise as a way to identify pairs of genes that, when mutated, may serve as biomarkers for the same treatment. These observations promise to be useful in guiding drug-repurposing efforts.

## Methods

### Somatic mutation data

In July 2016, we downloaded all available TCGA somatic mutation data via the National Cancer Institute’s Genomic Data Commons[11]. These data had been generated using high-throughput, exome-sequencing technologies. Using the *MuTect* tool[12], these data had been compared against germline variation using human reference genome *GRCh38* to make somatic calls. Subsequently, the variants had been annotated with the *Variant Effect Predictor* (VEP) tool[13] to record population frequencies and to predict effects of the mutations on gene function. We used these annotations to filter the somatic variant data. We excluded variants that did not pass MuTect’s quality-control criteria or that had a minor allele frequency greater than 0.01 in the ExAC database. Any mutation predicted by VEP to have a “LOW” or “MODIFIER” impact on protein function was excluded. We retained variants that had been predicted by SIFT[14] to be “deleterious” or “deleterious (low confidence) or” by Polyphen-2[15] to be “probably damaging” or “possibly damaging”. Although some of these variants were likely false positives, we preferred to err on the side of inclusion rather than exclusion, to minimize the chances of false negatives. In addition, we retained variants that contained no predictions for SIFT or Polyphen-2.

### Copy-number variation data

We downloaded copy-number variation (CNV) data that had been preprocessed and stored in the University of California Santa Cruz Xena database[16]. These data had been produced using whole-genome microarrays. The CNVs had been called using the GISTIC2.0 algorithm[17] and summarized to gene-level values. In addition, the gene-level values were thresholded to estimate whether each sample carried a homozygous deletion, a single-copy deletion, a low-level amplification, or a high-level amplification. We considered tumors with either a homozygous deletion or a high-level amplification to be “mutated”. We considered the remaining samples either to have been in a normal state or to have exerted only a modest effect, if any, on tumor transcription.

Finally, we aggregated the somatic mutation and CNV data to identify tumor samples that carried at least one genomic aberration in a given gene. We merged the somatic-mutation and CNV data to a single value per gene and tumor sample, using Boolean values to indicate whether the tumor harbored an aberration in a given gene.

### RNA-sequencing data

We downloaded gene-level, RNA-Sequencing data that had been preprocessed and aligned using the Rsubread package[18, 19] and had been summarized using the transcripts per million (TPM) method. For tumor samples that had been sequenced multiple times, we averaged expression values across these samples. We log2-transformed the RNA-Sequencing values to mitigate the effects of extremely high expression values and to enable easier visualization.

### Disease-drug-gene mapping data

We downloaded disease-drug-gene mappings that had been curated via a crowdsourcing effort and had been made available via the CIViC database[7]. We used the October 1, 2016 version of this database and focused solely on evidence where the relationship between gene and drug was “Supported”.

### Machine-learning analysis

Our analysis focused exclusively on TCGA samples for which data were available for all three molecular types (somatic mutation, CNV, and RNA-Sequencing). To reduce noise and computational complexity, we limited the data to 325 genes considered to play a role in canonical cancer pathways, according to the *Pathways in Cancer* diagram in the Kyoto Encyclopedia of Genes and Genomes (KEGG) [20].

Using the genome-aberration data, we categorized each tumor according to cancer type and whether an aberration had been identified in a given gene. We generated a training set by identifying samples that had at least one mutation in one of the frequently mutated genes (frequency threshold varied; see Results). We then excluded tumor samples that had a mutation in more than one of these genes. We used the resulting data set for a classification analysis, using genes as class labels. The remaining samples were set aside as a “test set”. As a way to assess our therapeutic predictions, we limited genes in the test set to those that were described in the CIViC database.

Using the training set, we performed 5-fold cross validation to evaluate our ability to predict mutation status for each gene. Later, we trained a model on the entire training set and made predictions for the test set. In both cases, we used the Random Forests classification algorithm[21] with default parameters, other than that we requested probabilistic predictions.

Because mutation frequencies varied considerably across genes, we subsampled the data. For example, if the minimum number of mutated samples across all selected genes were 15, we would randomly select 15 samples for each gene. We repeated this process 10 times and averaged the results across the various subsampled results. This approach ensured that class imbalance would not bias our results.

### Analysis pipeline

We wrote scripts in the *Python* programming language[22] to parse, filter, and summarize the input data. To enable easier analyses in subsequent steps, we restructured the data into “tidy data” format[23]. In performing the analysis steps and producing graphics, we used the *R* programming language[24]. These steps were aided by the following packages: *readr*, *dplyr*, *magrittr*, *ggplot2*, *RColorBrewer*, *randomForest*, *mlr, coin*, and *AUC*[25–33]. The entire pipeline executes in 5-15 minutes on a laptop computer with 4 cores and 16 GB RAM. We placed all analysis code and the tidy data in an open-access repository at https://osf.io/ndjkg.

To evaluate differences in expression for individual genes, we calculated p-values using Student’s t-test and then performed a Bonferroni correction to account for all possible gene pairs (n = 52,650) in our data.

### Results

We sought to identify genes whose transcriptional profiles were similar to each other when mutated in tumors. From TCGA, we obtained somatic-mutation data for 10,391 tumor samples. The filtering steps (see Methods) reduced the number of somatic mutations by 93.5%, mostly due to the removal of synonymous variants and common variants. Within our cancer-related genes of interest (n=325), we observed a total of 45,950 somatic mutations (5.57 per sample). We observed 52,012 high-level amplifications (8.63 per sample) and 19,037 large-scale, homozygous deletions (4.13 per sample) within these genes. After we removed samples that lacked data for at least one type of aberration, data for 9,300 patients across 25 distinct cancer types remained.

As might be expected, strong, tissue-specific patterns were apparent in the gene-expression data (Fig 1A). In particular, expression levels for Acute Myeloid Leukemia (LAML) and Lower Grade Glioma (LGG) were clearly distinct from the other cancer types, likely in part because these cancers affect myeloid blood cells and glioma cells, respectively, whereas most other cancer types affect epithelial cells. Although in some contexts, these tissue-specific differences may provide valuable insights about how tumor biology differs among cell types, our preliminary analyses revealed that these differences caused a strongly confounding effect that would make it difficult to derive pan-cancer insights. Therefore, to correct for these effects, we applied the *ComBat* algorithm[34], using cancer type as batch. This adjustment reduced cancer-type effects (Fig 1B). We used these corrected data in our analyses whenever we aggregated data across multiple cancer types.

**Figure 1.**
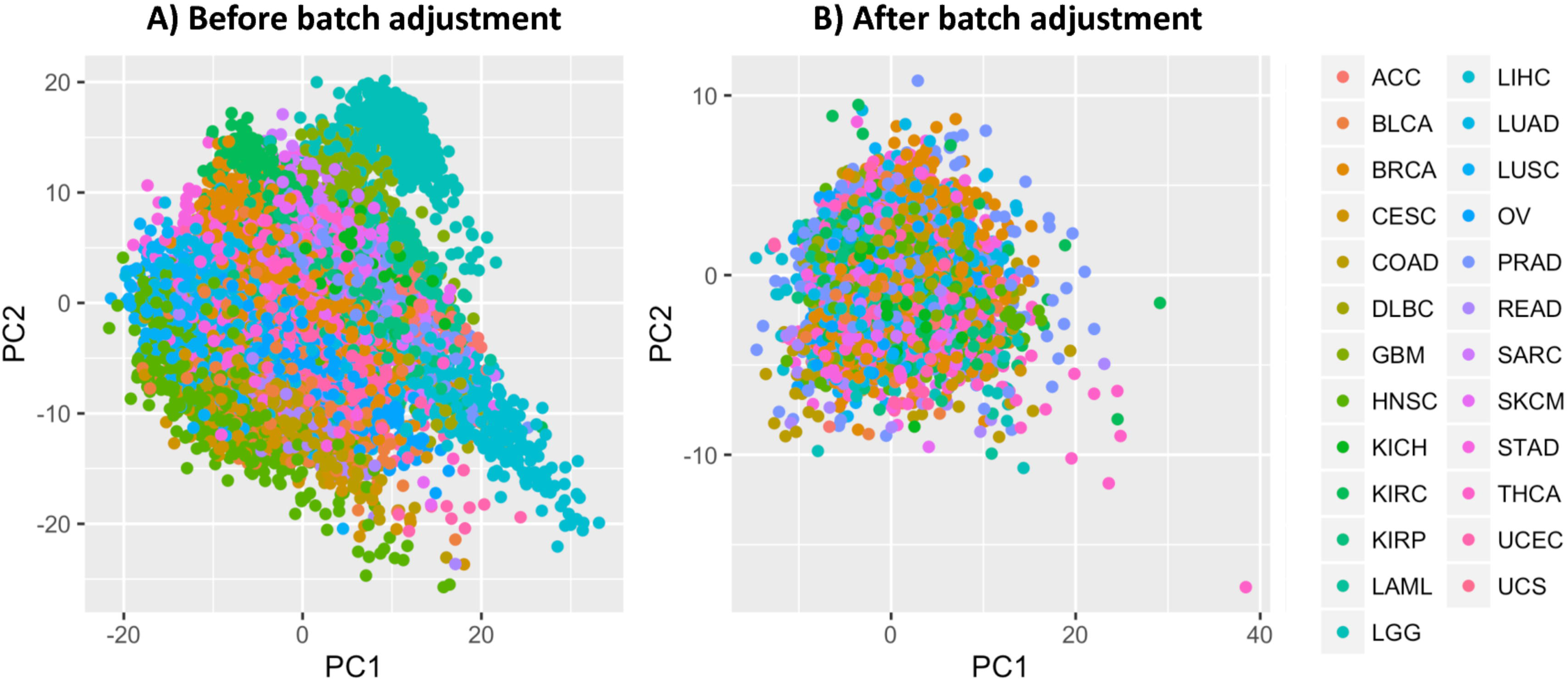
Principal component plots illustrating the relationship between cancer type and gene-expression levels. Values for the first two principal components are shown. Each point represents data across all genes for a given tumor sample. The colors and legend indicate the cancer type of each tumor sample. A) Distinct, tissue-specific patterns were present in the original data. B) After a regression-based adjustment for cell-type effects, these patterns were no longer prominent in the data. Cancer type abbreviations are listed on the TCGA web site.

### Single-gene analysis

For the first analysis phase, we examined the relationship between genomic aberrations and expression levels of individual genes. In lung adenocarcinomas (n=515), the most frequently mutated genes were TP53 (n=233), CDKN2A (n=109), KRAS (n=95), CDKN2B (n=91), and EGFR (n=82). For decades, mutations in KRAS and EGFR have been recognized to play a critical role in lung adenocarcinoma and have been used to classify these tumors into subtypes based on mutation status[35]. Codons 12, 13, or 61 are typically mutated in KRAS [36]; similar mutations affect homologous genes HRAS and NRAS, though infrequently for lung adenocarcinomas. EGFR is often affected by point mutations and amplifications. Due to its role as a cell-surface receptor, EGFR is the target of antibody-based therapies, including *Gefitinib*[37], and vaccine-based immunotherapies[38]. We examined the relationship between mutations and expression levels of these genes. Tumors that harbored an EGFR aberration—a mutation or a copy-number variation—expressed EGFR at significantly higher levels than tumors that harbored a KRAS aberration but no EGFR aberration (Fig 2A). In addition, tumors with a KRAS aberration expressed KRAS at significantly higher levels than samples with an EGFR aberration but no KRAS aberration (Fig 2B). Although genomic aberrations often do not correlate strongly with gene-expression levels of the affected gene, we observed strong patterns for these genes. EGFR interacts with KRAS indirectly, via the Ras-Raf-MEK-ERK signaling cascade; thus EGFR aberrations may have led to suppression of KRAS activity in some samples. Indeed, mutations in these genes are often mutually exclusive, and KRAS mutations have been associated with a lack of sensitivity to EGFR-targeted therapies, such as *Gefitinib* and *Erlotinib*[39]. These observations suggest that, despite some likely overlap, the downstream effects of aberrations in these genes are distinct. In contrast, we examined differences in CDKN2A expression between samples that harbored either an EGFR or a KRAS aberration. No significant difference in expression was observed (Fig 2C), as might be expected, given that CDKN2A is far downstream from these genes. We observed similar patterns after merging data across all 25 cancer types (Fig 3).

**Figure 2.**
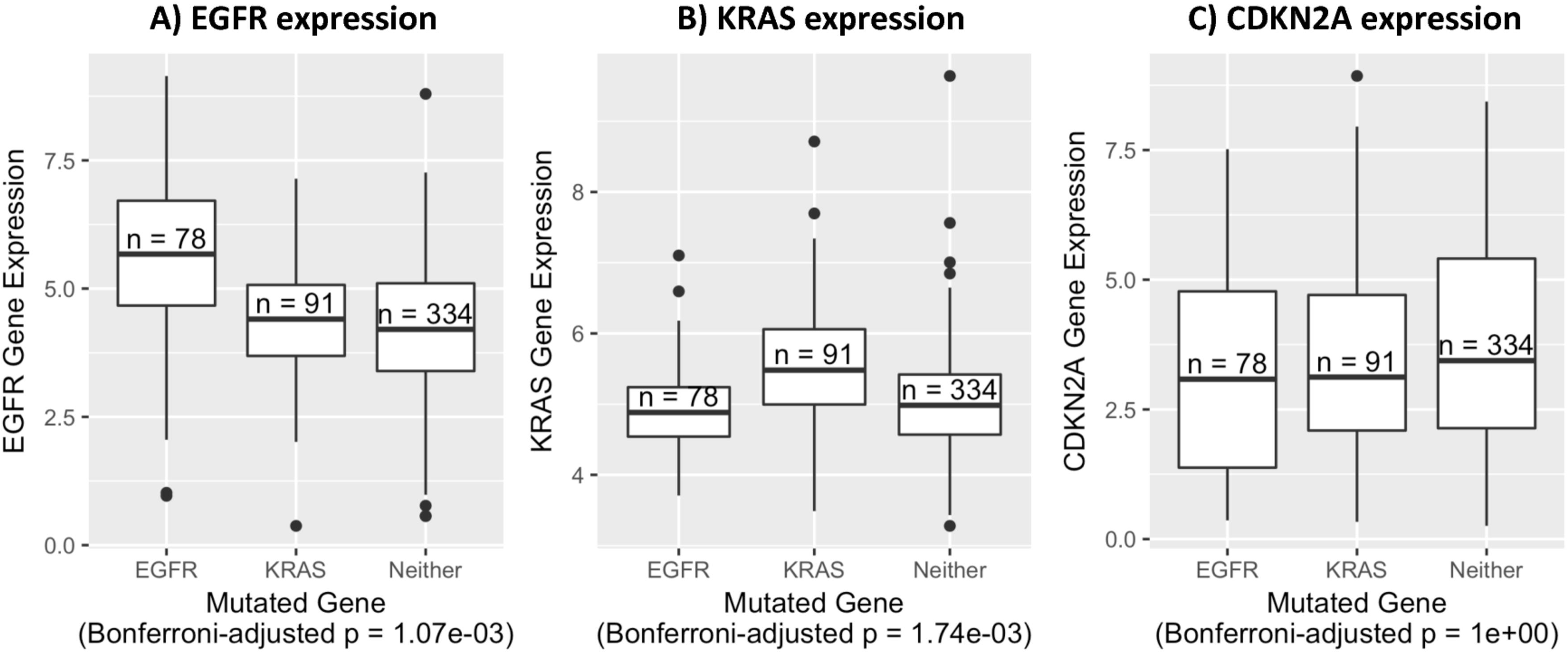
Associations between genomic aberration status and expression levels of individual genes in lung adenocarcinoma samples. A) EGFR expression levels for tumors with an aberration in EGFR, KRAS, or neither of these genes. B) KRAS expression levels for the same tumors. C) CDKN2A expression levels for the same tumors. Expression levels are log-transformed, transcripts-per-million values. P-values were calculated using Welch’s t-test and adjusted using a Bonferroni correction.

**Figure 3.**
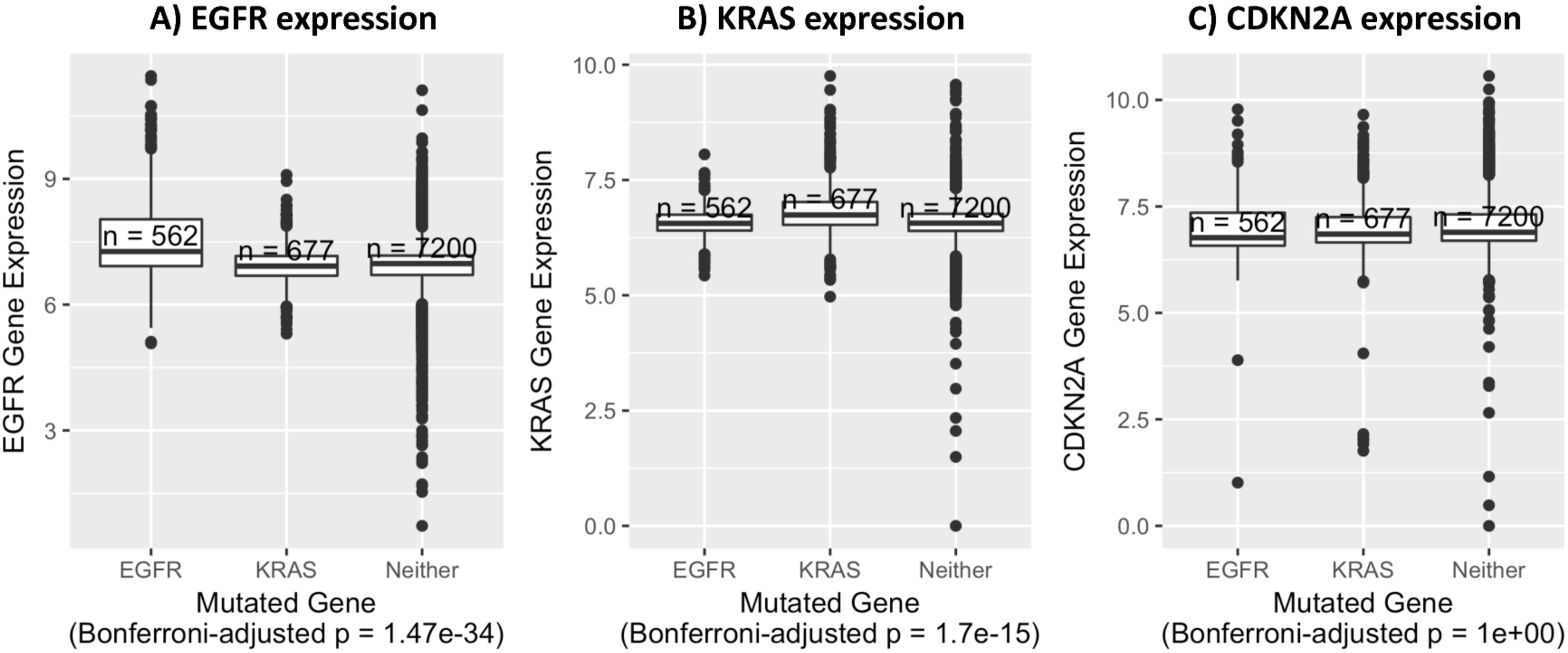
Associations between genomic aberration status and expression levels of individual genes across 25 cancer types. A) EGFR expression levels for tumors with either an aberration in EGFR, KRAS, or neither of these genes. B) KRAS expression levels for the same tumors. C) CDKN2A expression levels for the same tumors. Expression levels are log-transformed, transcripts-per-million values. P-values were calculated using Welch s t-test and adjusted using a Bonferroni correction.

### Supervised-learning analysis

Although it is interesting to examine the effects of genomic aberrations on individual genes, most aberrations cause transcriptional responses across a broad range of genes expressed throughout tumor cells. To account for these broad effects, we used a supervised-learning approach (see Methods). Initially for lung adenocarcinomas, we identified genes that were mutated in at least 10 tumor samples. Then we excluded tumor samples that had a mutation in more than one of these genes. Finally, we retained genes that still had mutations in at least 10 tumor samples, resulting in a set of tumors with mutually exclusive mutations. We used cross validation to evaluate the ability of the Random Forests classification algorithm to predict gene-mutation status based on the gene-expression levels. We interpreted relatively high accuracy levels to mean that the downstream, transcriptional effects of mutations in these genes were distinct from each other. Fig 4 illustrates predictions for the five genes selected. Across these genes, the average classification accuracy was 0.42—whereas 0.20 would be expected by chance in this scenario. For each gene, we calculated the area under the receiver operating characteristic curve (AUROC), an alternative measure of classification accuracy that accounts for the probabilistic nature of predictions. The highest AUROC was 0.81 for EGFR (0.50 expected by chance). The lowest AUROC was 0.58 for KRAS. These findings suggest that EGFR mutations have a particularly distinct effect upon transcription levels in lung adenocarcinomas. To get a sense for similarities and differences among genes, we calculated the Spearman correlation coefficient for probabilistic predictions between each pair of genes. We observed no strong positive correlations for lung adenocarcinoma, providing evidence that the transcriptional effects of mutations in KRAS, EGFR, MET, NTRK1, and PIK3CA are distinct from each other (Fig 5). We saw a modest negative correlation (rho = -0.18) between EGFR and KRAS—consistent with our single-gene analysis—and a strong negative correlation (rho = -0.47) between EGFR and MET. Both EGFR and MET are from the receptor tyrosine kinase family, encode for cell-membrane receptors, and act as oncogenes when mutated. Although crosstalk mediated by microRNAs may occur between these proteins [40], the negative correlation we observed may indicate that these proteins cause very different downstream effects and thus should be addressed differently from a therapeutic standpoint. Indeed, it has been shown that MET amplification causes resistance to *Gefinitib*, an EGFR inhibitor[41].

**Figure 4.**
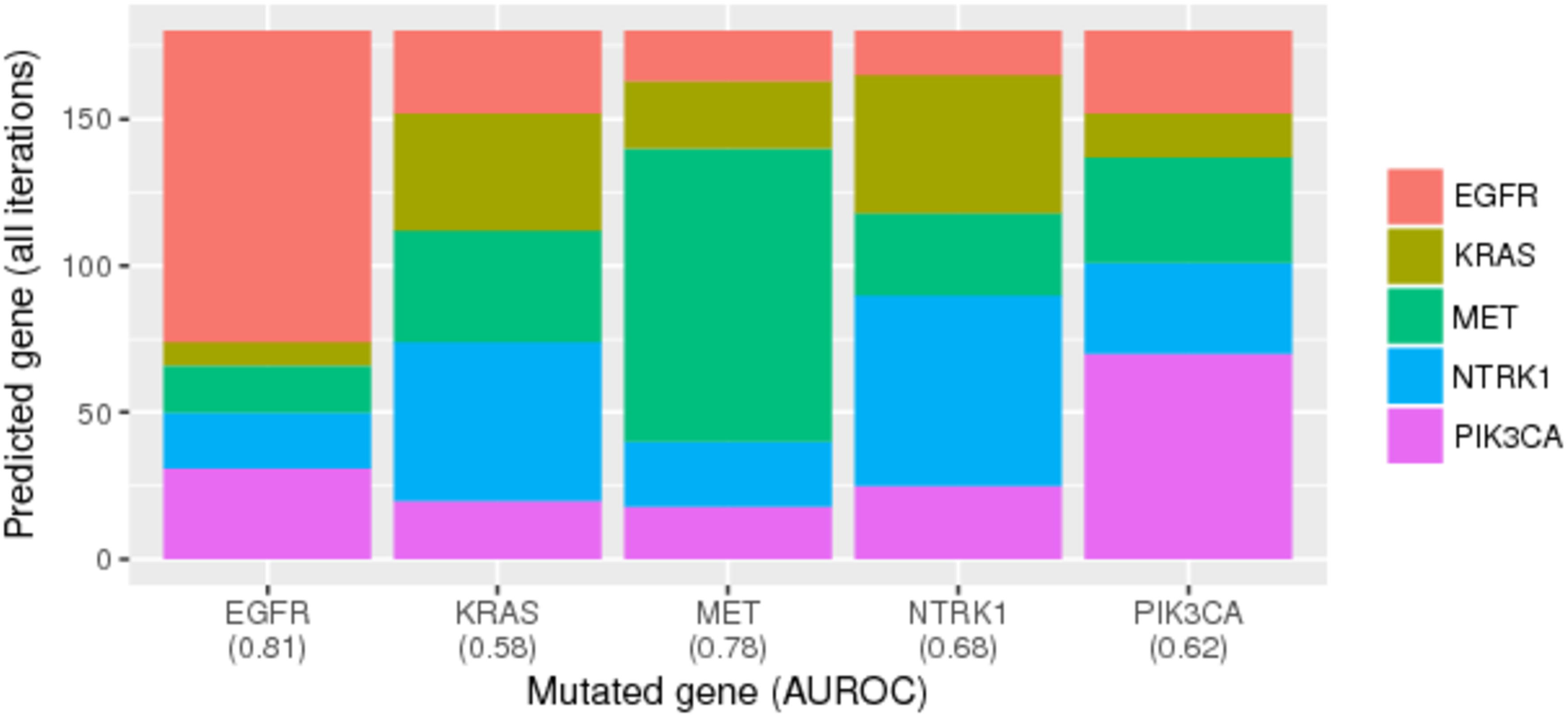
Random Forest predictions, trained on gene-expression data, of gene-mutation status for lung adenocarcinoma. We identified genes that had been mutated in at least ten tumor samples and that contained no mutation in other genes that had been mutated in at least ten samples. Then using cross validation, we evaluated how well the Random Forest classification algorithm could identify which gene was mutated in a given sample. The algorithm produced a probabilistic prediction for each gene (class), and we evaluated these predictions using the area under the receiver operating characteristic curve (AUROC). The x-axis labels indicate AUROC values for each gene. Relatively high AUROC values indicate that gene-expression levels are highly predictive of gene-mutation status and thus suggest that mutations in these genes exert a characteristic effect on gene-expression levels.

**Figure 5.**
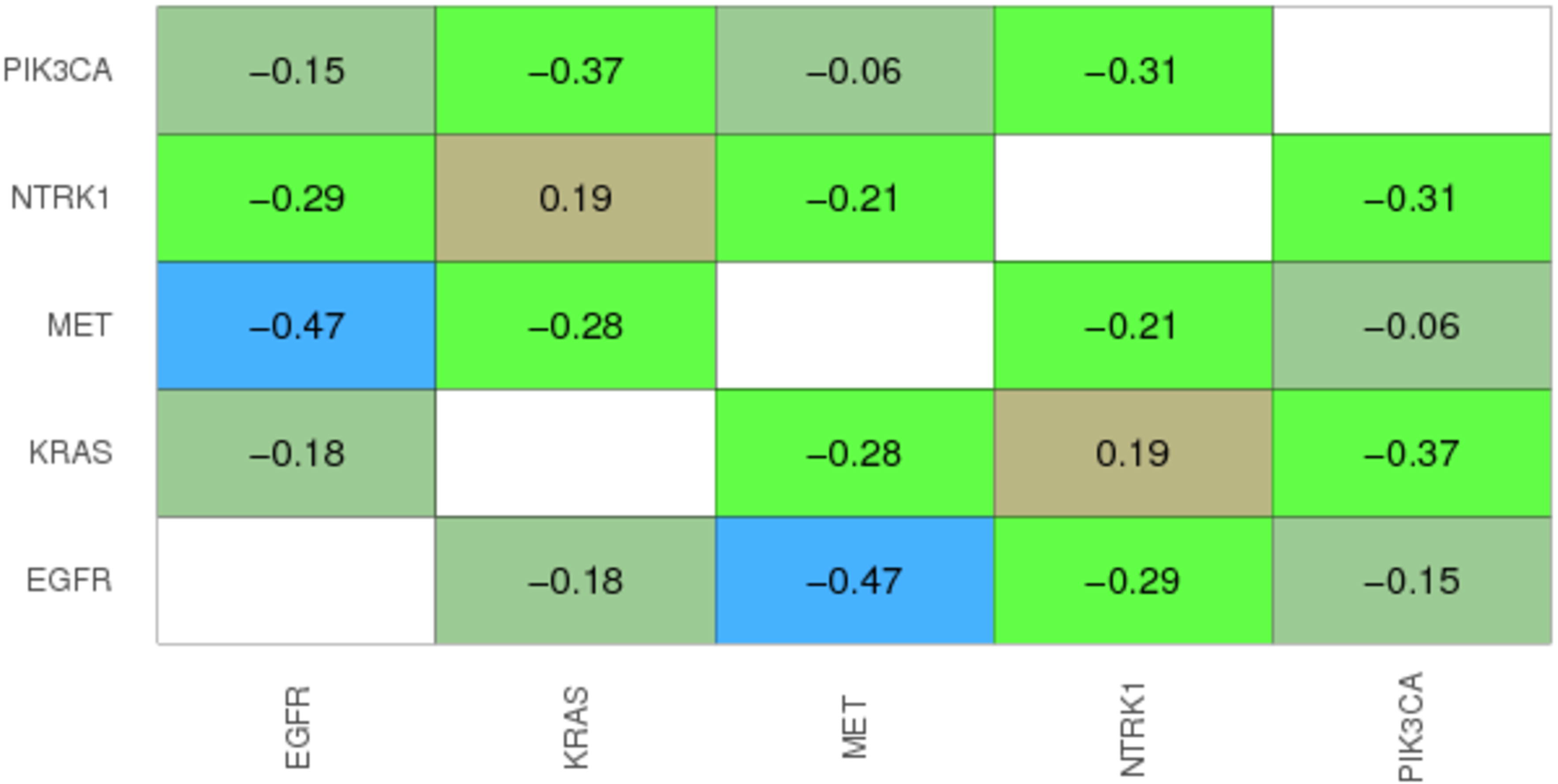
Heatmap showing Spearman correlation coefficients between gene pairs for lung adenocarcinomas. The data values and colors represent correlation coefficients between probabilistic predictions for pairs of genes that exhibited mutually exclusive mutations. Warmer colors represent higher levels of correlation, while cooler colors represent lower levels of correlation.

We applied the same methodology to breast tumors. Seven genes passed the threshold for inclusion in the training set, and the average accuracy was again 0.42 (0.14 expected by chance). The genes with the highest individual AUROC were BIRC5 and TP53. Both play a role in tumor cells ability to evade apoptosis, but our results suggest that mutations in these genes operate via distinct mechanisms. The strongest positive correlation (rho = 0.49; see Figure S1) was between PTEN and PIK3CA, two genes known to play important roles in breast tumorigenesis (Fig 6). PIK3CA is an oncogene, activated by receptor tyrosine kinases and plays an important role in cellular proliferation via the PKB/Akt signaling cascade. PTEN tightly regulates PIK3CA; thus, PKB/Akt activity is expected to increase when PTEN loss (via mutation or deletion) has occurred[42]. Our results coincide with the expectation that the downstream effects of genomic aberrations in these genes are similar. Indeed, aberrations in these genes have been shown to correlate similarly with prognostic factors[43].

**Figure 6.**
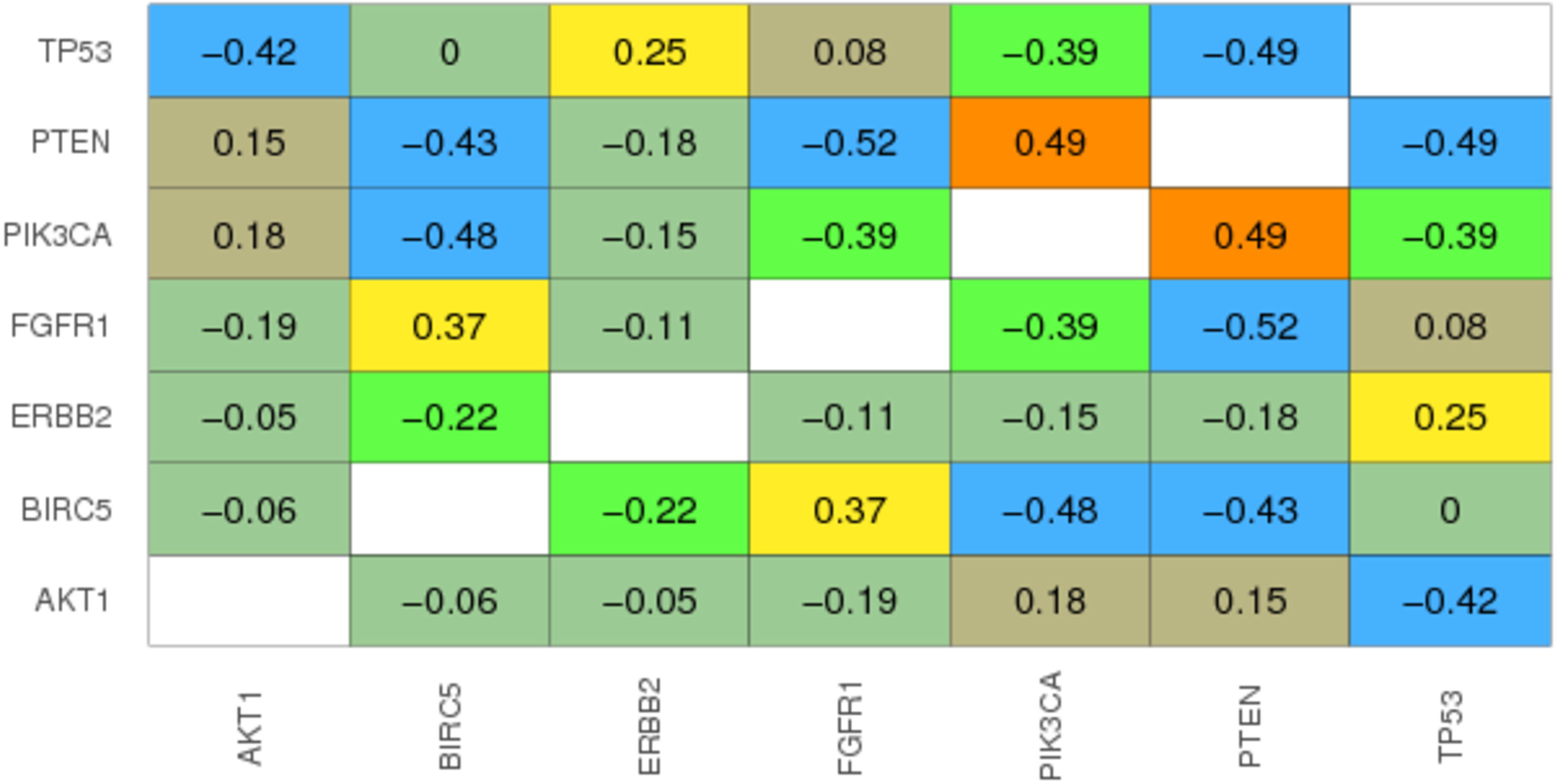
Heatmap showing Spearman correlation coefficients between gene pairs for breast carcinomas. The data values and colors represent correlation coefficients between probabilistic predictions for pairs of genes that exhibited mutually exclusive mutations. Warmer colors represent higher levels of correlation, while cooler colors represent lower levels of correlation.

We repeated this process for other cancer types—bladder carcinoma, head/neck squamous carcinoma, ovarian cystadenocarcinoma, metastatic skin melanoma, stomach adenocarcinoma—that had enough data; we required that at least three genes be mutated at least ten times in a mutually exclusive manner. *Additional files 1-8* indicate correlation coefficients, nominal p-values, and Bonferroni-adjusted p-values for each pairwise comparison. The strongest negative correlation (rho = 0.83; see Figure S2) was between FGFR1 and FGFR3 in bladder carcinoma. Experimental work has demonstrated that both genes are activated via mutations and the genes play distinct roles in regulating bladder-tumor growth[44]. The genes that showed the most significant differences in expression between FGFR1- and FGFR3-mutated tumors were SMAD3, FGFR3, FN1, LAMA1, and FGFR1 (Figures S3-S7). In addition to growth signaling, these genes play roles in regulating extracellular matrix adhesion and intracellular signaling.

The correlations we observed between pairs of mutated genes were often tissue specific; for example, a strong relationship between PTEN and PIK3CA was observed in breast carcinomas but not in head/neck squamous carcinomas, stomach adenocarcinomas, or ovarian cystadenocarcinomas.

Next we extended the same methodology to all 25 cancer types in a pan-cancer approach. We increased the threshold to 50 for the minimum number of mutually exclusive mutations per gene. Nine genes passed this threshold (see Fig 7). Perhaps due to the larger sample sizes per gene, accuracy increased to 0.50 (0.11 expected by chance). The AUROC was extremely high for BRAF (0.97) and VHL (0.93), suggesting that the transcriptional effects of aberrations in these genes are particularly distinct. However, mutation status was predicted accurately for all nine genes, with a minimum AUROC of 0.73 (APC gene). Several interesting relationships between genes were apparent, including a strong correlation (rho = 0.44) between TP53 and RB1, which play critical roles in regulation of the cell cycle and DNA repair[45, 46]. FGFR1 was strongly correlated (0.42) with PIK3CA, likely due to FGFR1’s role as an upstream activator of PIK3CA. Perhaps surprisingly, FGFR1 predictions were also correlated strongly with RB1 predictions; however, it has been shown recently that CDK6, a key regulator of RB1, is regulated by MIR9, a microRNA that also regulates FGFR1 activity[47].

**Figure 7.**
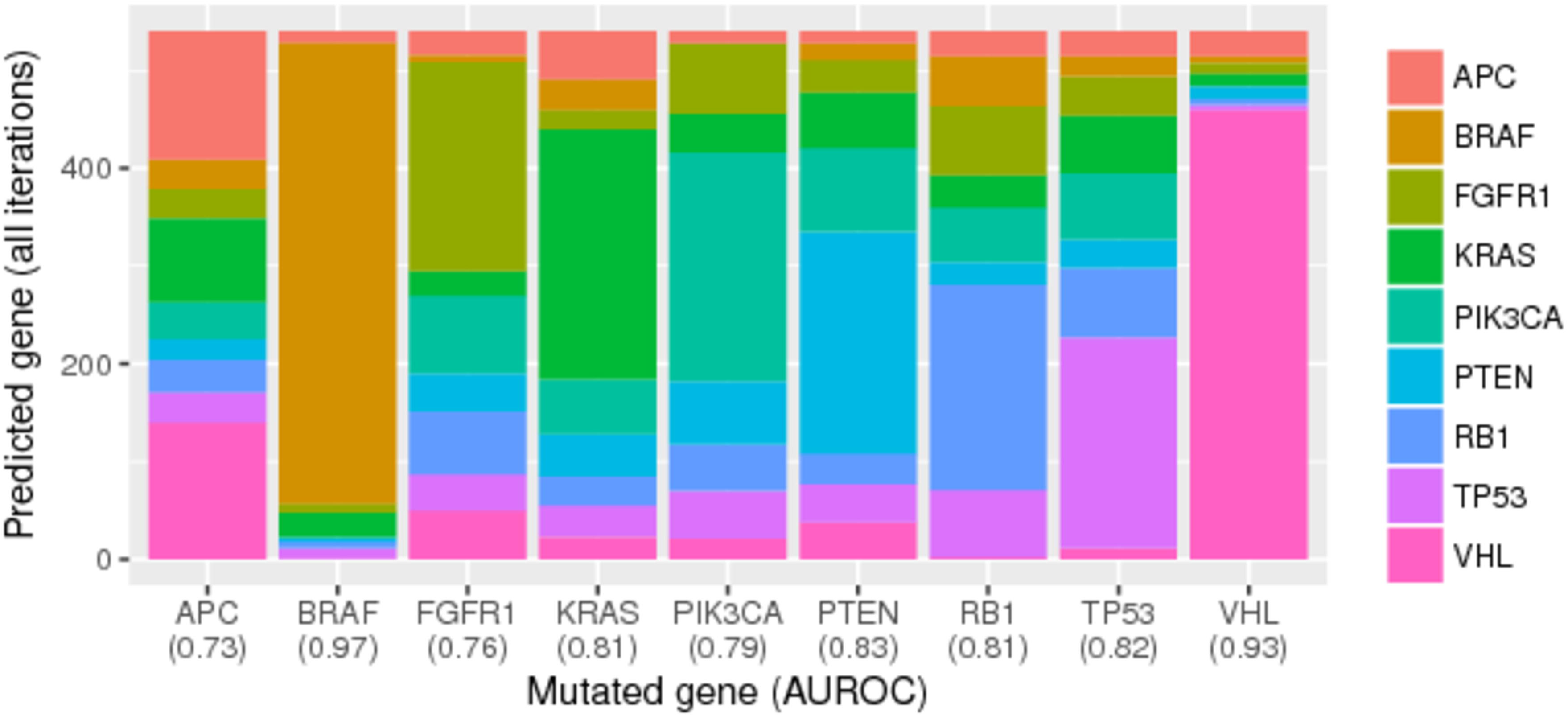
Random Forest predictions of gene-mutation status across 25 cancer types. We identified genes that had been mutated in at least 50 tumor samples and that contained no mutation in other genes that had been mutated in at least 50 samples. Then using cross validation, we evaluated how well the Random Forest classification algorithm could identify which gene was mutated in a given sample. The algorithm produced a probabilistic prediction for each gene (class), and we evaluated these predictions using the area under the receiver operating characteristic curve (AUROC). The x-axis labels indicate AUROC values for each gene. Relatively high AUROC values indicate that gene-expression levels are highly predictive of gene-mutation status and thussuggest that mutations in these genes exert a characteristic effect on gene-expression levels.

For the last stage of our analysis, we trained the Random Forests classification algorithm on the full training set and made predictions for tumor samples in the test set. We interpreted that probabilistic predictions coinciding with the actual mutation status of genes in the test set would indicate that the expression profiles of genes in the training set were similar to the expression profiles of genes from the test set. We hoped this analysis would provide insights about genes that are mutated rarely. Using a minimum threshold of five mutated samples (test set) and focusing initially on breast cancer, we found that MMP2 predictions were strongly associated with predictions for PIK3CA (0.47), PTEN (0.49), and AKT1 (0.28), which operate via the same pathway. Like MMP2, PIK3CA and AKT1 play important roles in cancer cell migration and metastasis[48–50].

We also observed a strong positive correlation (0.33) between TP53 and AKT2 and a modest correlation between ERBB2 and AKT2 (0.20). AKT2 mediates TP53 activity via MDM2[51]. *Trastuzumab* was originally developed as a targeted therapy for ERBB2 (Her2) amplification; more recently, aberrant AKT2 expression has been associated with longer time to progression and overall patient survival in Her2-positive patients[52]. We also observed a slightly negative correlation (-0.20) between AKT1 and AKT2 predictions; although the proteins encoded by these genes operate within the same signaling cascade, their roles in regulating cell migration and differentiation are distinct[48].

When we made test-set predictions for all 25 cancer types, we noted a few modestly positive correlations. The first was between VHL and MTOR1. VHL inactivation leads to constitutive activation of HIF-2 and/or HIF-1. In clear-cell renal carcinomas, the downstream effects of HIF activation are inhibition of *mTor Complex 1*[53]. The second positive correlation was between NRAS and BRAF. These genes interact directly with each other and operate via the Ras → Raf → MEK → ERK cascade. Indeed, the CIViC database indicates that several antibody-based treatments—including *Cetuximab*, *Selumetinib*, and *Vemurafenib*otarget tumors with mutations in either of these genes.

Although we used the ComBat software to correct for tissue-type effects and used a principal component analysis to visually verify the effects of this correction (Figure 1), we conducted a follow-up evaluation to assess whether tissue specificity might still have influenced our results, because, in many cases, mutations occur in a tissue-specific manner[54]. Initially, we used the Random Forest algorithm to predict tumor type based on the expression data that had not been adjusted using ComBat. Then we repeated this process using the ComBat-adjusted data. Using this approach, we could predict tumor type with >90% accuracy for both versions of the expression data. Even though ComBat had adjusted for tissue specificity, a subtle footprint remained, which the Random Forest algorithm was able to detect. In a second follow-up evaluation, we used the Random Forest algorithm to predict gene-mutation status using only tumor type (instead of gene-expression data). Although tumor type predicted mutation status less accurately than the gene-expression data (47% and 50%, respectively), these results confirm that tumor type is confounded with mutation status and thus that our pan-cancer resultsoand results from other pan-cancer studies that examine the relationship between genomic and transcriptomic variation—should be interpreted with caution. However, it is difficult to distinguish between correlation and causation; certain tissue environments (driven by gene expression) may select for somatic variants in certain genes, and/or certain somatic variants may strongly influence the tissue specificity of cancer.

## Discussion

We have developed a computational approach that uses publicly available, molecular-profiling data to identify genes that have similar (or different) effects on gene expression in human tumors. Our overarching goal was to develop a methodology that can be used to guide drug-repurposing efforts and, more generally, to help cancer researchers make sense of the vast complexity and heterogeneity of tumors. Using data from The Cancer Genome Atlas, we observed relationships that recapitulate what was previously known about canonical cancer pathways and treatment responses. Alternative methods have primarily evaluated molecular data in a low-throughput manner, have examined one type of molecule at a time, or have considered the expression of individual genes associated with mutations; in contrast, our method accounts for broad-ranging effects of genomic aberrations on gene expression within cells. Using a supervised-learning approach, we found that we could predict mutation status, often with high accuracy. This provides evidence that many mutations confer a clear and distinct effect on transcriptional responses within downstream genes.

To increase interpretability, we made simplifying assumptions and simplified our approach in several ways. We limited our analysis to genes known to play a role in cancer, as described in KEGG’s *Pathways in Cancer* diagram. We assumed that the effects of genomic aberrations are mutually exclusive. Even though this assumption may not hold in every case, it reflects current approaches that are used to prioritize targeted cancer therapies. For example, even though it is clearly understood that tumors with mutations in EGFR harbor many variants in genes other than EGFR, mutations in EGFR are used as a biomarker for treating lung adenocarcinomas with therapies such as *Gefitinib* and *Erlotinib*. In addition, evidence suggests that EGFR and KRAS mutations occur in a mutually exclusive manner and that tumors with KRAS mutations fail to respond to these drugs[39]. Such findings suggest that mutations in individual genes can strongly influence treatment responses, despite a background of other mutations. To an extent, our goal was to identify such scenarios.

In addition, we ignored the potential impact of epigenomic factors, such as DNA methylation and miRNA expression, as well as gene fusions. Our approach could be refined in future studies to use such observations to indicate whether a given gene is “mutated”. Although genomic aberrations are often thought to be the main drivers of tumorigenesis, epigenomic factors often play a critical role in modulating tumor activity and/or interacting with genomic aberrations. For example, DNA hypermethylation of promoter regions can cause transcriptional silencing of tumor-suppressor genes; BRCA1 hypermethylation has been shown to alter responses to platinum-salt therapies[55]. miRNAs can also play important roles in regulating tumor transcription[56]; in chronic lymphocytic leukemia, two miRNAs on human chromosome 13q14 occur frequently [57] and likely play important roles in regulating tumorigenesis and treatment responses.

We treated all mutations equally within a given gene, regardless of genomic loci or mutation type; but in many cases, there may be considerable heterogeneity across genomic loci and mutation types. As sample sizes increase over time, it will be more feasible to sub-classify mutations in finer detail.

We focused primarily on the genome- and cancer-related aspects of this work rather than on algorithmic aspects. In future work, it would be valuable to compare (and perhaps combine) multiple algorithms as a way to optimize our approach. We used the Random Forests algorithm because it has been shown to deal effectively with high-throughput transcriptomic data, executes quickly, and is more amenable to *post hoc* interpretation than many other algorithms[58].

Finally, although we have demonstrated a potential to learn about the effects of rare variants, our analysis touched only briefly on such variants. To evaluate the reliability of our method, we focused on genes that had been affected by at least a modest number of mutations. However, we believe this methodology can be applied in cases where a given mutation has been observed in only a single tumor, potentially providing insights for “n-of-1” clinical trials.

## Conclusions

We have used supervised-machine learning to integrate genomic and transcriptomic data across 9300 tumors and 25 cancer types to aid in deciphering the downstream effects of genomic aberrations on tumor transcription. This approach has potential to guide development of treatment biomarkers and to understand similarities and differences among genes that play a role in specific types of cancer and across multiple cancer types. We hope this approach will be useful to pharmacologists and clinical trialists who seek to identify relationships between genomic aberrations and treatment responses. In particular, we hope this methodology will reduce barriers for drug-repurposing efforts so that existing treatments can be used on tumors with no current treatment biomarker. We believe such approaches will reduce the costs of developing new cancer drugs and increase the number of tumors that can be treated in a targeted manner.

### Ethics approval and consent to participate

The Brigham Young University Institutional Review Board granted an exemption for this study (#E14522). This study uses only publicly available data.

## Declarations

### Availability of data and material

All tidy-data files, analysis code, and analysis outputs (including some figures not included in this paper) are publicly accessible at https://osf.io/ndjkg. We encourage others to extend and refine our methods.

## Authors’ contributions

SRP conceived and designed the study. Both authors developed the analytical pipeline. JBD parsed, filtered, and tidied the data. SRP performed the machine-learning analysis. JBD prepared and organized the code and data. Both authors wrote the paper and prepared figures for the paper. Both authors read and approved the final manuscript.

## Acknowledgements

We thank T. James Lee for providing insights on processing data from TCGA. We thank the Fulton Supercomputing Lab at Brigham Young University for providing computational resources that enabled us to complete this study. We thank the patients who donated tumor samples to TCGA Consortium and consented to share the resulting data with the research community.

## Additional files

***Additional file 1***

*Multi-class Correlations − BLCA* (.tsv). Correlation coefficients and measures of significance for each gene pair in our analysis of bladder carcinoma.

***Additional file 2***

*Multi-class Correlations − BRCA* (.tsv). Correlation coefficients and measures of significance for each gene pair in our analysis of breast carcinoma.

***Additional file 3***

*Multi-class Correlations − HNSC* (.tsv). Correlation coefficients and measures of significance for each gene pair in our analysis of head/neck squamous carcinoma.

***Additional file 4***

*Multi-class Correlations − LUAD* (.tsv). Correlation coefficients and measures of significance for each gene pair in our analysis of lung adenocarcinoma.

***Additional file 5***

*Multi-class Correlations − OV* (.tsv). Correlation coefficients and measures of significance for each gene pair in our analysis of ovarian cystadenocarcinoma.

***Additional file 6***

*Multi-class Correlations − SKCM* (.tsv). Correlation coefficients and measures of significance for each gene pair in our analysis of metastatic skin melanoma.

***Additional file 7***

*Multi-class Correlations − STAD* (.tsv). Correlation coefficients and measures of significance for each gene pair in our analysis of stomach adenocarcinoma.

***Additional file 8***

*Multi-class Correlations − All Cancer Types* (.tsv). Correlation coefficients and measures of significance for each gene pair in our pan-cancer analysis.

